# Aging as a defense strategy against parasites

**DOI:** 10.1101/143362

**Authors:** Richard Y. Chin

## Abstract

The teleology of aging has been one of the more vexing and controversial question in biology. One potential evolutionary driver of programmed aging is selection pressure from parasites and other infectious organisms. While selection pressure from parasites and other infectious organisms have long been considered by many biologists to have led to the evolution of sexual reproduction, it has only rarely been considered as a potential driver for evolution of aging, a biological process that likely evolved contemporaneously with sexual reproduction. Here I describe stochastic simulations of host and parasite populations with senescence as an independent variable. The results show that populations with more rapid senescence bear lower parasite loads and oscillate more quickly through alternate phenotypes with differential resistance against parasites. I conclude that programmed aging and death may promote host evasion of parasites in a co-evolutionary competition against parasites.

## Introduction

The most common theories regarding aging can be divided into two groups: programmed aging theories and non-programmed aging theories.(1) Programmed aging theories, which historically have been less favored than non-programmed aging theories by most evolutionary biologists, posit that aging is an adaptive process that grants species that age an evolutionary advantage over species that do not. One example of such theory is Weismann’s theory that as organisms age, they accumulate mutations and somatic that reduce their fitness and that culling such individuals from the population can improve the overall fitness of the population.(2)

Non-programmed aging theories postulate that aging is a degenerative process that do not confer an evolutionary advantage. In fact, one of the more widely held non-programmed aging theories argues that post reproductive age, the selection pressures against deleterious genes disappears, and those genes accumulate in the genome, as opposed to deleterious genes that manifest prior to the reproductive age, which are subject to purifying selective pressure.(3)

Whether aging is programmed or non-programmed is important. If aging is a programmed process rather than a non-programmed degenerative process., then there are significant implications for the direction of anti-aging research and furthermore, aging may be a more tractable disease than previously believed, because it may mean that aging can be prevented or delayed by re-programming the senescence genes.

Most of the non-programmed theories are not fully convincing. For example, if the span of reproductive age determines onset of senescence, it begs the question of what drives reproductive age to cease. Furthermore, in species such as humans where males continue to be fertile for much longer than females, sometimes until death, it doesn’t explain why women don’t age more rapidly than men. Nonetheless, non-programmed aging theories have generally been more convincing that the programmed aging theories, and have been more favored by aging theorists.

However, recently, this prevailing view of aging as a non-programmed, degenerative process has been challenged by new discoveries. For example, recent discovery of genes that appear to influence the rate of aging have led to the recognition that at least in some cases, genes can have a profound effect on the rate of senescence. Furthermore, it appears that the senescence-related genes are sometimes conserved over widely disparate organisms. For example, IGFR-1 gene and its homologs appear to control the rate of aging across multiple species including *C. elegans*, bats, dogs, and perhaps even humans.(4) In *C. elegans*, mutation of *daf-2*, a member of the IGF-1 receptor family, increases lifespan by 100%.(5) Similarly, long-lived bat species, such as Brandt’s bats that live for 40 years, carry a mutation in the IGF-1 receptor gene that confers longevity.(6) Similarly, small dog species, which live for 15 to 20 years vs. 7 to 10 years typically seen with large dog species, all carry a mutation in their GFR-1 receptor gene.(7) Small dogs are in addition virtually immune to cancer.(8) Likewise in humans, Laron dwarfism is caused by a mutation in the growth hormone receptor gene, which controls IGF-1 levels, and those with the dwarfism also are virtually immune to diseases such as cancer and diabetes, although evidence of increased longevity is not clear.(9)

Furthermore, recent demonstration that transfer of blood from young animals to old animals can reverse some features of aging, as well as a report that transfer of microbiome from young killifish to old killifish can substantially increase the old killifish’s lifespan have all suggested that aging may be a programmed, rather than a degenerative process.(10),(11),(12)

However, a convincing explanation for programmed aging has been lacking. One of the weaknesses in the previously suggested theories such as Weismann’s is that the selection pressure would have be strong enough to cause virtually all organisms to develop aging. Theories that posit accumulation of mutations over a lifetime as driver of senescence, and other similar theories have not held up to scrutiny.(1)

One potential source of evolutionary pressure for aging are parasites and other infectious organisms. Many of the most sophisticated biological processes, such as restriction enzyme, adaptive immune system, and CRISPR system, are result of evolutionary pressure from infectious organisms. Furthermore, a leading explanation for one of the most ubiquitous biological phenomenon other than aging, sexual reproduction, is parasite resistance. Infectious organisms make up one of the strongest drivers of evolution.

Only rarely, however, has infectious organisms been considered as a potential explanation for evolution of programmed aging.(13) One previously proposed theory is that senescence serves either to reduce population density or to reduce effective population density. The reduction of effective population density is through increase in host genetic diversity. The reduction in population density inhibits disease transmission and reduces the likelihood of disease-mediated species extinction. Otherwise, there has been little to no recognition that a factor that might have driven the evolution of sexual reproduction, infectious organisms, might play a role in the evolution of aging.

In fact, sexual reproduction and aging are tightly coupled. In semelparious organisms, obligatory and immediate senescence is usually coupled to reproduction. The prototypical example is salmon, which age and die in the process of reproduction. The death of animals such as salmon post reproduction is believed by some biologists to be the result of the energy expenditure required to generate eggs milt, and not a programmed process, but several lines of evidence militate against that. First, salmon undergo changes typically associated with aging, albeit at a very rapid pace, during the reproductive process, including formation of Alzheimer’s disease-like plaques, and atherosclerosis-like lesions.(14) Second, removal of endocrine organs delays onset of aging and death, even if the salmon has invested energy in migration and egg production. Third, Atlantic salmon undergo similar strenuous migration and spawning as Pacific salmon but many species are iteroparous.(15)

In fact, in many semelparous organisms, aging can be delayed. For example, in octopus, the female normally dies within ten days of reproduction. But removal of glands called optic glands prevents aging and death for several months. In annual plants, which normally flower only once in their lives, if the seed is prevented from forming after fertilization, the plant will flower repeatedly.(15)

The reverse can also be true, In many organism, senescence can be triggered by reproduction. For example, in some *C. elegans*, females respond to mating by reducing their lifespan by 40%. (16),(17) Initially, this was throught to be perhaps secondary to the energy required to produce eggs, but subsequently it was demonstrated that mating without egg production could trigger senescence, as could exposure to chemical secreted by males.(18),(19) Similarly, in seed beetles, males appear to secrete chemicals that control the longevity of females.(20) In drosophila, in contrast, exposure to pheromones from females, mediated by taste receptors on males’ forelegs, appear to reduce lifespan in males under certain circumstances.(21)

Furthermore, given that aging is generally, although not exclusively, limited to multicellular sexually reproducing organisms, it is likely that the two traits evolved at the same time. Furthermore, it is reasonable to hypothesize that the evolutionary pressures that led to sex also may be involved in aging.

According to the Red Queen hypothesis, sex evolved as a mechanism for periodically changing host defense mechanisms against parasites, in the face of parasites’ adaptability to those defense mechanisms.(22) There is supportive evidence that strains of parasites and host susceptibility to the strains may cycle in a periodic fashion.(23),(24) If oscillatory evolution is a key constituent of successful arms race against parasites, the oscillatory period may be an important variable in the strategy. The faster the host can cycle through the repertoire of defensive genetic combinations, more easily it may be to evade the parasite. It stands to reason that rapid senescence post reproduction may accelerate the cycling of the host parasite resistance phenotypes.

I therefore postulated that parasite defense may be a reasonable teleological explanation of programmed aging. Programmed senescence may be an important variable in the effectiveness of the sex-based parasite defense system, because the periodicity of the defense cycles and the effectiveness of the cycling in reducing parasite burden may be influenced by the lifespan of the hosts. Specifically, the faster the host populations can change the distribution of resistance genes, more successfully they may evade the parasites.

In order to test the hypothesis, I created a model of host and parasite populations. The model assumed a fixed host population, with three phenotypes of hosts and three phenotypes of parasites. Each host phenotype was completely resistant to infection against one parasite phenotype, partially resistant to a second parasite phenotype, and not resistant to one phenotype in a cyclic fashion, such that host phenotype A was completely resistant to parasite phenotype A and partially resistant to parasite phenotype C; host phenotype B was completely resistant to parasite phenotype B and partially resistant to parasite phenotype A; and so on. This assumption is consistent with a previously described cyclic parasite resistance pattern.(23)

Each host was assumed to be at risk for dying each year, based on a base rate of non-senescence-related mortality and after reaching a predefined age of senescence, a rate of senescence-related rate of death based on an inverse of age onset of senescence. Each host that died was replace by a new host, whose phenotype was based on the population distribution of the host phenotypes in the population during the prior year, modified by reproductive penalty for hosts that were infected, and modified by a fixed regeneration factor that simulated recombination due to sexual reproduction. The regeneration factor allowed for regeneration of phenotypes even when they have disappeared from the population, consistent with the theory proposed by Hamilton that alleles with transient low fitness is stored and not eliminated, and re-expressed at a subsequent time.(22) Each uninfected host could be infected by a parasite, determined by a transmission rate. The phenotype of the parasite for the infection was determined by the population distribution of parasite phenotypes during the prior year, modified by a fixed regeneration factor that simulates recombination due to sexual reproduction. If the host was resistant to the parasite phenotype, then the likelihood of infection was modified by the resistance factor. Each host was permitted to be infected by one parasite strain at a time.

## Materials and methods

The simulation was programmed in Chipmunk Basic 3.6.7b6 for Mac OS X. The model assumed a fixed population of 100 organisms, with three phenotypes of hosts and three phenotypes of parasites. Each iteration of the simulation was for 500 years. There were total of 1,000 iterations of the simulation performed for each lifespan. Each set of iterations was performed for onset of senescence set at 2, 5, 10, and 20 years.

The initial host phenotypes were randomly assigned with equal probability across each of the three phenotypes, and the initial parasite burden level was set at 40%, with parasite phenotypes were randomly assigned with equal probability across each of the three phenotypes. Initial age was randomly assigned with equal probability for each age between 0 and onset of senescence.

Each host phenotype was 100% resistant to infection against one parasite phenotype, 90% resistant to a second parasite phenotype, and not resistant to one parasite phenotype in a cyclic fashion, such that host phenotype A was completely resistant to parasite phenotype A and partially resistant to parasite phenotype C; host phenotype B was completely resistant to parasite phenotype B and partially resistant to parasite phenotype A; and so on.

Each host was assumed to be at 5% risk for dying each year, based on a base rate of non-senescence mortality and after reaching a predefined age of senescence, plus an additional probability death based on an inverse of age of senescence. For instance, for 2-year onset of senescence population, each organism had 50% senescence-based risk of dying each year after year 2.

Each host that died was replaced by a new host, whose phenotype was based on the population distribution of the phenotypes in the population the prior year, modified by 50% reproductive penalty for hosts that were infected, and modified by a fixed regeneration factor that regenerated each phenotype at a fixed rate of 0.33% percent per year.

Each uninfected host was subject to a probability of infection by a parasite each year, determined by the proportion of infected hosts during the previous year multiplied by 1.2. Each host that was infected was randomly assigned a parasite phenotype, based on the proportion of parasite phenotype in the population the previous year, modified by 3.33% fixed risk of infection by each parasite phenotype each year regardless of the parasite population the previous year. If the host was infected by a parasite phenotype to which it was resistant, the host rejected the infection based on the resistance factor. Each host was permitted to be infected by up to one parasite strain at a time.

The mean parasite load was calculated by summing the mean proportion of hosts with an infection for each simulation and dividing by the total number of simulations.

## Results

All populations cycled through the host and parasite genotypes over time. The periodicity of the oscillations in the 2-year onset of senescence simulation was the shortest, and the periodicity of the oscillations in the 20-year onset of senescence simulation was the longest. (Figs 1a and 1b). The 2-year senescence onset population cycled through median of 29 genotypes and the 20-year senescence population cycled through a median of 7 genotypes.

**Fig 1a and 1b.**
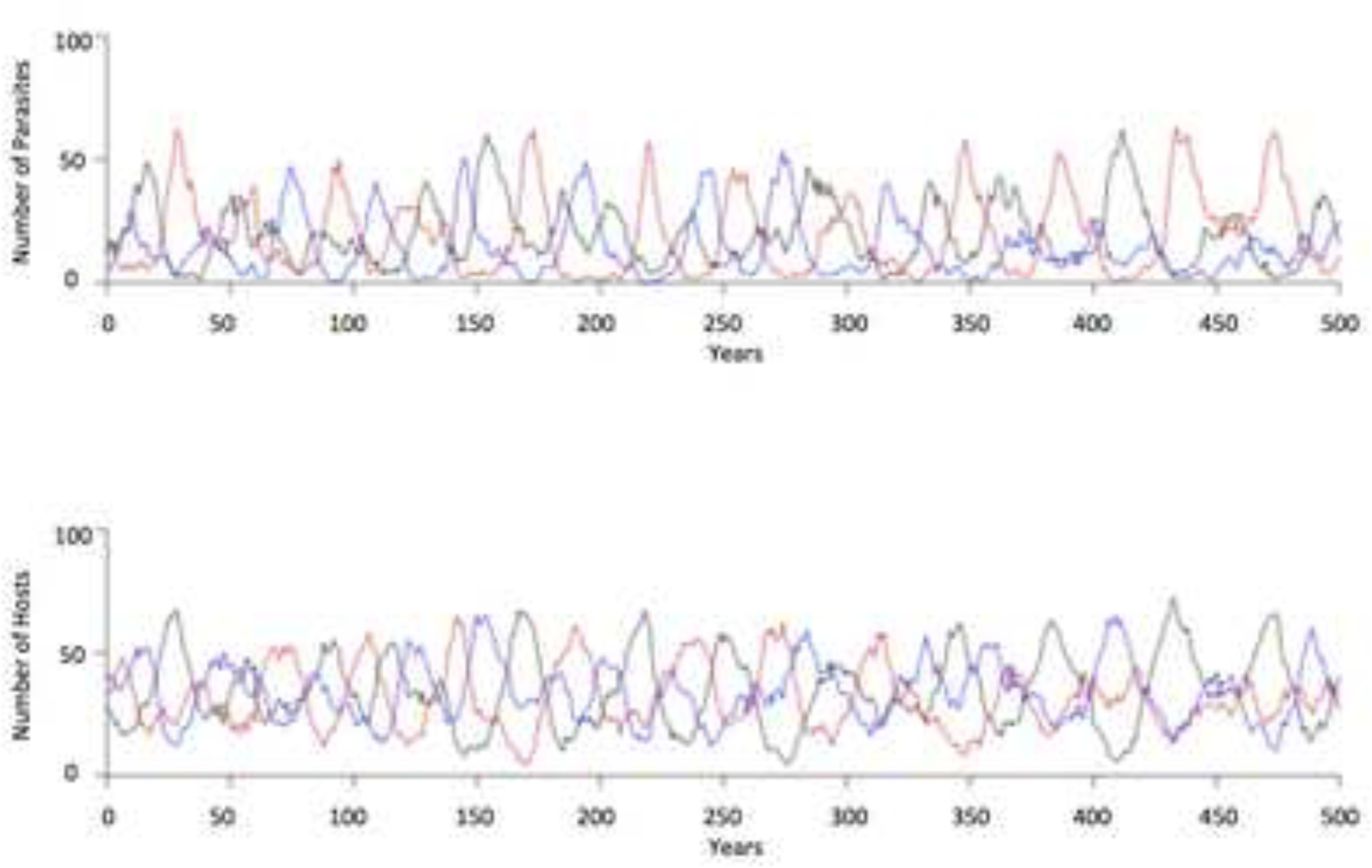
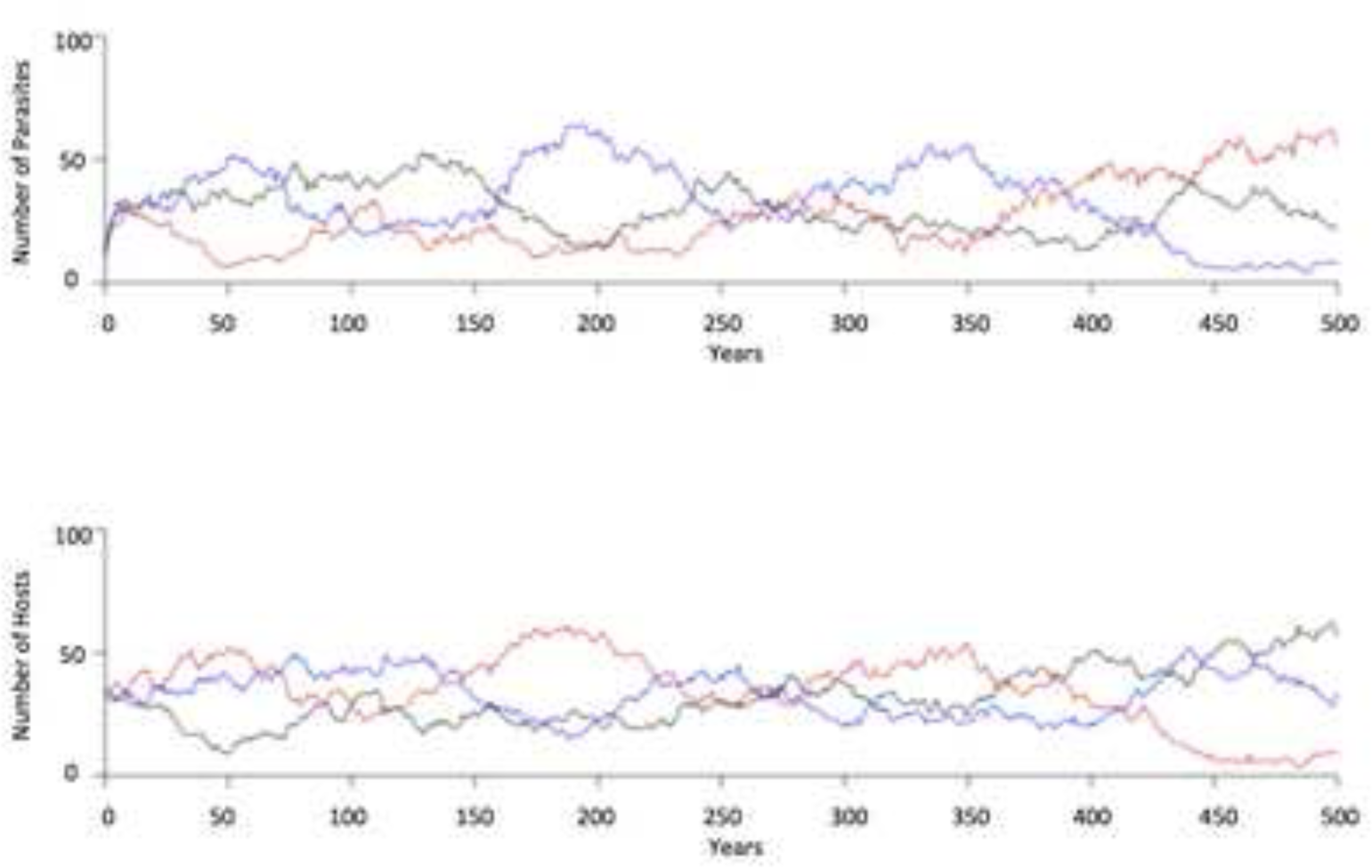
Representative stochastic simulations of host and parasite populations. **a.** Simulation with 2-year onset of senescence. The top graph is the parasite population, and the bottom graph is the host population. Vertical axis is the number of organisms, and the horizontal axis is the years. Each color represents one of three phenotypes. The corresponding color on the graphs represents one parasite phenotype and the host phenotype that is resistant to the parasite phenotype. **b.** Simulation with 20-year onset of senescence.

Table 1 illustrates mean parasite load after 1,000 iterations of the simulation for the 2-year, the 5-year, the 10-year, and the 20-year senescence onset populations. As hypothesized, the parasite load was correlated with lifespans, with populations with shorter onset of senescence exhibiting lower overall parasite burdens than populations with longer onset of senescence. The differences were statistically significant, with p-value <0.0001 by one-way ANOVA.

**Table 1.** Average Parasite Load.

|  | 2-Year | 5-Year | 10-Year | 20-Year |
| --- | --- | --- | --- | --- |
| Average Parasite Load | 59.20% | 80.93% | 87.62% | 90.39% |

The mean parasite load, as measured by percentage of hosts that are infected in each year for each of the simulations, after 1,000 iterations of 500-year simulations.

Unlike in a previously reported model, senescence did not have a significant effect on average host genetic diversity and because it was a fixed population size model, the population size did not change.

## Discussion

I performed simulations to test whether the onset of senescence could influence the cycling periodicity and the average parasite load in a host:parasite population model.

The results from this model suggests that under a certain set of assumptions, decreased lifespan can result in lower parasite load, along with more rapid oscillation of host and parasite resistance genotypes. The decrease in oscillatory period and the decrease in parasite load may have a causal relationship, and support the hypothesis that if senescence is an adaptive, programmed trait, then evolutionary pressure from parasites may be a factor driving its pervasiveness, much as it may have been the driving factor for the evolution of sexual reproduction. This is consistent with the fact that in order for sexual reproduction to lead to effective parasite evasion, the parasite resistance genotype distribution in the host population must evolve as rapidly or more rapidly than the genotype distribution of parasites.

## Conclusions

This study is the first to show that programmed senescence can increase the frequency of host:parasite genotype cycling and lower average parasite load in a host population. If sexual reproduction evolved as a method to cycle between alternate parasite resistance genotypes at a population level, then programmed aging may act to multiply the effectiveness of the cycling. Aging and reproduction are tightly linked in many species, and since the evolution of both traits likely occurred contemporaneously, it stands to reason that both traits evolved in response to the same evolutionary pressure, and that aging may enhance the effectiveness of the sexual reproduction strategy.

